# A lipid-driven, microbe-independent mechanism of acne via *Lrig1^+^* follicular progenitor cells

**DOI:** 10.1101/2025.08.30.673221

**Authors:** Takashi Sugihira, Masakazu Tamai, Kohei Takagi, Barbora Salcman, Tetsuro Kobayashi, Alison J Hobro, Seitaro Nakagawa, Yukiko Takeuchi, Eri Ishikawa, Motoko Maekawa, Yuji Owada, Naohiro Inohara, Hiroki Takahashi, Nicholas I Smith, Sho Yamasaki, Manabu Fujimoto, Yuumi Nakamura

## Abstract

Long-chain saturated fatty acids (lcSFAs) are abundant in the skin, but their pathogenic roles in acne remain unclear. In human sebum profiling, C16:0, the most abundant lcSFA, was the only fatty acid significantly elevated in acne and correlated with inflammatory comedone counts. We then established a mouse model that faithfully recapitulates human acne phenotypes, in which topical C16:0 penetrated the epidermis and induced sebocyte hyperplasia, comedogenesis, and follicular inflammation. Mechanistically, C16:0 activated keratinization and inflammatory pathways and drove sebocyte-lineage differentiation from *Lrig1^+^*progenitor cells, which function as cutaneous lipid sensors. These effects persisted in germ-free mice and were unaffected by fatty acid transporter modulation, while *MyD88* signaling was partially required for inflammatory cell recruitment. Together, our findings identified C16:0 as a human acne-associated lipid that recapitulates disease hallmarks through a microbe-independent, lipid-driven pathway, highlighting *Lrig1^+^*cells as central hubs in remodeling of the pilosebaceous unit.

## Introduction

Acne vulgaris (acne) is the most prevalent inflammatory skin disease, affecting up to 85% of adolescents and often persisting into adulthood.^1^ The condition typically begins during puberty, when rising androgen levels stimulate sebaceous gland hyperactivity, a hallmark of acne pathophysiology.^1^

Sebum secreted from sebaceous glands is a complex lipid mixture composed of squalene, glycerol esters, wax esters, cholesterol, and free fatty acids (FFAs).^2,3^ The rate of sebum excretion correlates with acne severity and is predictive of clinical outcomes.^1^ Accordingly, sebum hypersecretion has long been considered the primary sebaceous abnormality in acne. However, acne pathogenesis involves more than excess sebum production. The mechanisms underlying follicular hyperkeratinization (comedogenesis) and subsequent inflammation remain poorly understood, despite evidence implicating microbial colonization.

Acne lesions originate within the pilosebaceous unit through a series of events, beginning with comedogenesis, abnormal differentiation and excessive cornification of the follicular infundibulum. This process leads to keratinocyte accumulation, ductal obstruction and ultimately follicular distension which promotes inflammation.^4^ The skin-resident bacterium *Cutibacterium acnes* metabolizes triglycerides into FFAs, and its density increases with higher sebum levels.^1,5^ Follicular rupture and bacterial exposure amplify the inflammatory cascade, culminating in pustules, nodules, and cysts, all indications of the severe end of the acne spectrum.

While FFAs have long been proposed to act as proinflammatory agents in acne, their exact pathogenic roles in vivo remain poorly defined. To date, animal modeling has been limited to rabbit ear comedogenesis assays using lauric acid (C12:0),^6^ a minor component comprising less than 1% of human skin FFAs.^7^ In contrast, the predominant FFAs in human sebum include C14:0, C15:0, C16:0, C16:1, 2-methyl-C17:0, and C18:1, while odd-chain FFAs are primarily derived from bacterial metabolism, including by *C. acnes*.^7,8^ Although a mouse model has been previously used to study epidermal keratinization via topical FFA applications (C16:0, C18:0, C16:1, C18:1), this study has primarily employed hairless mice (HR-1) with no analysis of the hair follicle compartment.^9^ Other models involving intradermal injection of *C. acnes*, with or without topical lipid application, have been used to evaluate inflammatory responses in the skin.^10,11^ However, these models do not allow for the specific assessment of follicular responses.

These observations prompted us to ask whether topical application of structurally distinct FFAs to mouse skin could recapitulate key features of human acne, including comedogenesis and follicular inflammation. Specifically, we investigated how differences in FFA chain length, whether derived from host sebum or microbial metabolism, influence acne-relevant pathological changes in the pilosebaceous unit.

## Results

### Host-derived saturated fatty acid C16:0 is the most abundant cutaneous lipid and correlates with inflammatory comedones

We recruited 50 Japanese females with either normal or acne-prone skin and quantified two types of facial lesions: inflammatory comedones (representing classical inflamed acne lesions) and non-inflammatory comedones (whiteheads and blackheads) (**Figure 1A and S1A**). To link lesion type with lipid composition, we analyzed sebum samples by gas chromatography-mass spectrometry (GC-MS). This revealed a spectrum of saturated FFAs (C12:0-C23:0) and monounsaturated FFAs (C16:1, C18:1) in the skin **(Figure 1B)**. Among these, C16:0 (palmitic acid) was the most abundant and the only one significantly elevated in acne-prone versus normal skin. In contrast, odd-chain FFAs such as C15:0 and C17:0, primarily produced by skin-resident bacteria, occurred at far lower levels and showed no enrichment in acne-prone individuals (**Figure 1B**). Quantitative analysis of host-derived even-chain FFAs further demonstrated that only C14:0 and C16:0 levels correlated positively with inflammatory comedone counts, whereas C12:0 and FFAs of C18:0 or longer showed no such correlation (**Figure 1C, 1D**, **S1B and Table S1**). Conversely, non-inflammatory comedones were significantly associated with longer-chain saturated FFAs ranging from C20:0 to C23:0 but not from C12:0 to C18:0 (**Figure 1D** and **S1B, Table S1**).

**Figure 1.**
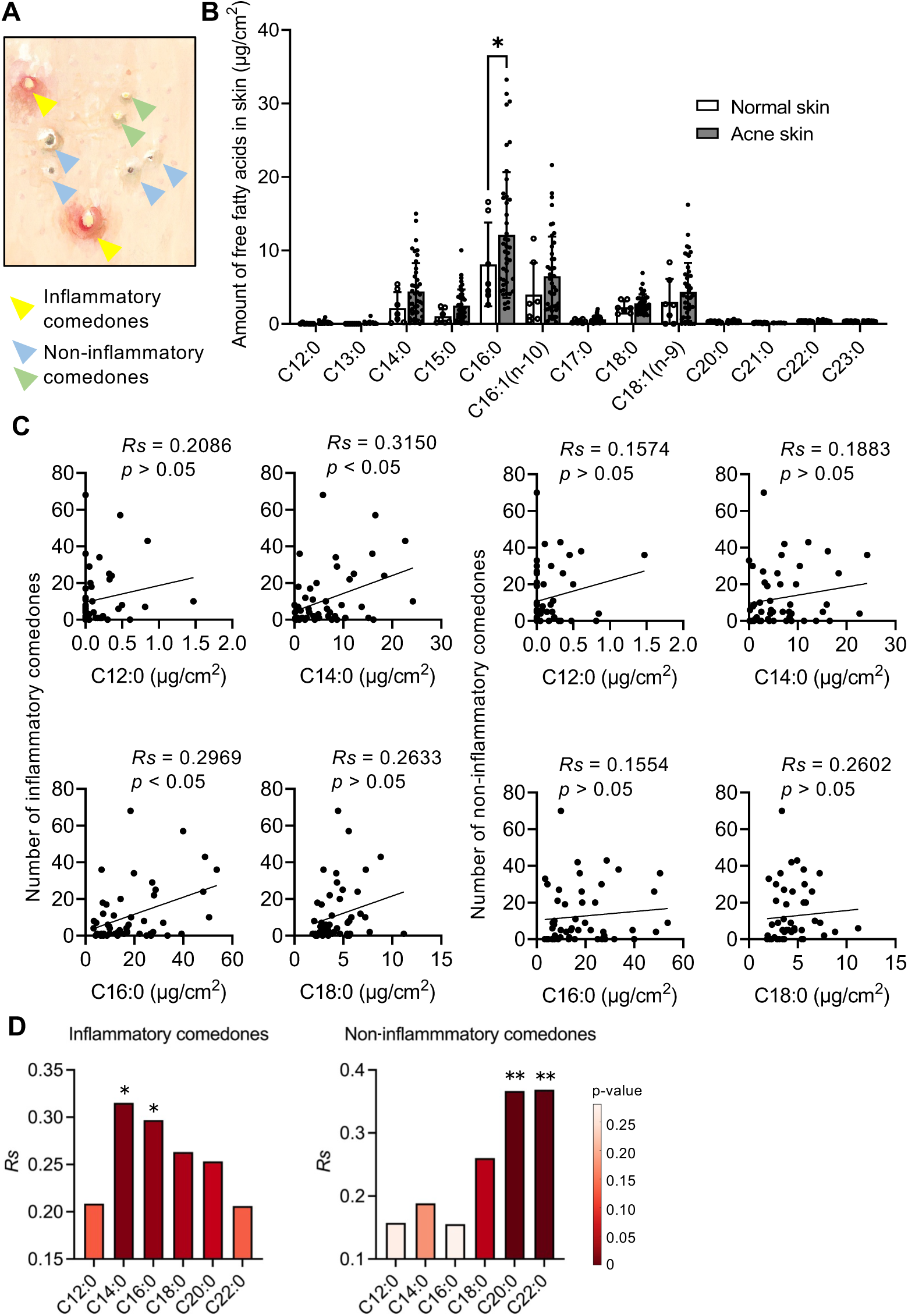
C16:0 is the dominant cutaneous saturated fatty acid and is associated with inflammatory acne. **(A)** Watercolor illustration depicts three major clinical types of comedones: inflammatory comedones (yellow arrows:erythematous papules with central follicular plugging) and non-inflammatory comedones (green arrows: blackheads, blue arrows: whiteheads). **(B)** Quantification of sebum-derived free fatty acids (FFAs) from the foreheads of individuals with acne-prone skin (*n* = 43) and healthy controls (*n* = 7), measured by GC-MS. Each dot represents an individual subject; bars indicate mean ± SEM. *p* < 0.05, Mann-Whitney test. **(C)** Correlation coefficients (*Rs*) between individual FFA levels and the number of inflammatory or non-inflammatory comedones. Each dot represents an individual subject. **(D)** Bars represent correlation coefficients (*Rs*) between the relative abundance of each FFA (C14:0, C16:0, C18:0, C20:0, and C22:0) and the number of comedones in each group (Inflammatory or Non-inflammatory). Bar color indicates the *p*-value for the correlation. Correlation coefficients (*Rs*) and *p* values were calculated using Spearman’s rank correlation test and are shown on each graph. ∗*p* < 0.05; ∗∗*p* < 0.01. Data in (B-D) are from one experiment obtained from two independent clinical studies.

Taken together, these findings indicate that C16:0 is a major host-derived lipid associated with inflammatory acne, independent of microbial FFAs. Furthermore, the data suggest that FFAs can be broadly categorized according to carbon chain length into those associated with classical inflammatory acne and those linked to follicular occlusion, such as whiteheads/blackheads.

### Comedogenesis is enhanced by longer saturated fatty acid chains, while C16:0 uniquely drives follicular inflammation

In vitro, FFAs from C14:0 to C18:0 induced proinflammatory chemokine expression (*CXCL1, CXCL2, Cxcl1, Cxcl2*) in primary human and mouse keratinocytes in a chain length-dependent manner, with C16:0 and C18:0 showing the strongest induction (**Figure 2A and 2B**). In mouse bone marrow derived macrophages (BMDMs), only C18:0 enhanced *Cxcl2* expression without affecting *Cxcr2* **(Figure 2C**).

**Figure 2.**
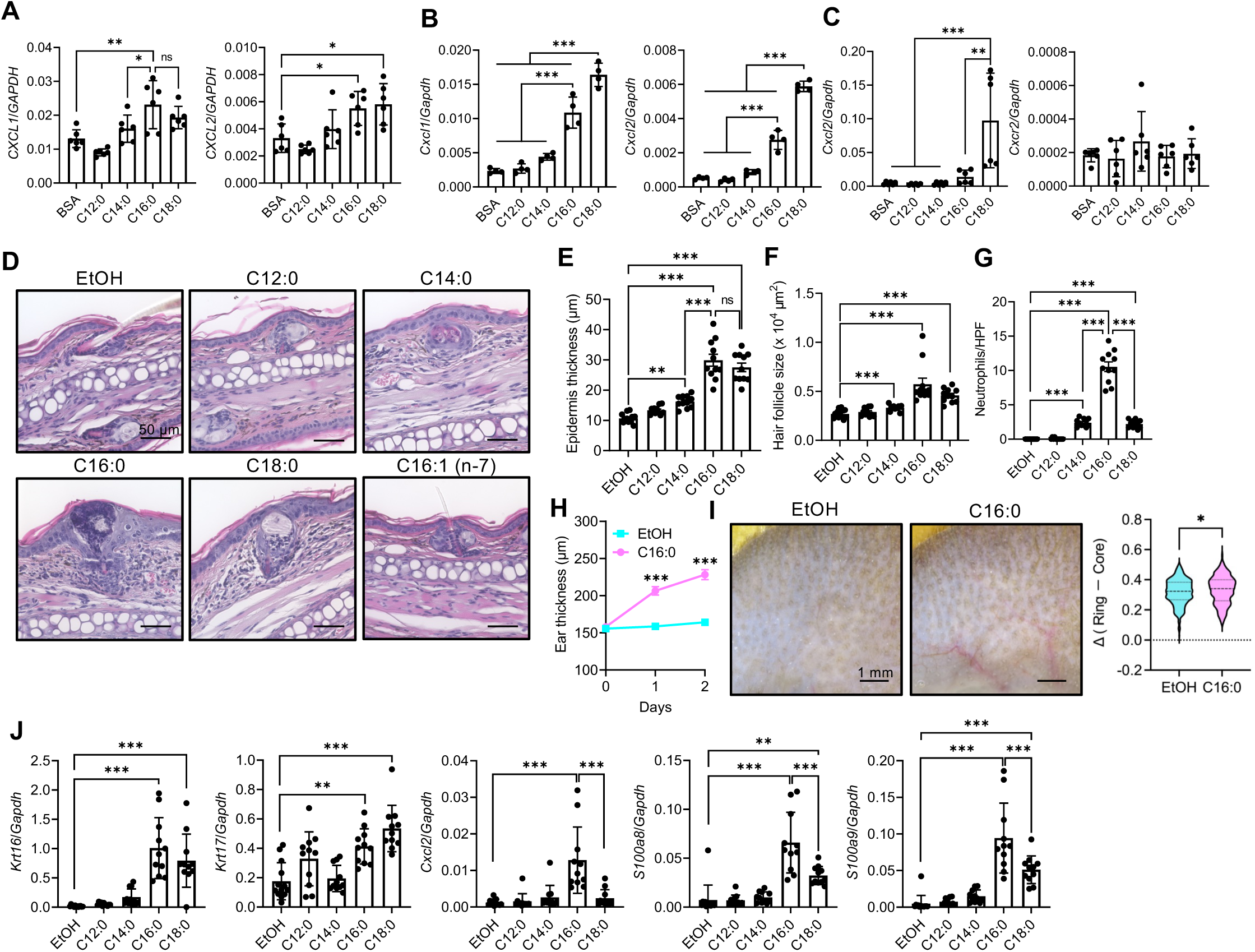
Comedogenesis is enhanced by longer saturated fatty acid chains, while C16:0 uniquely drives follicular inflammation. **(A, B)** Gene expression of proinflammatory chemokines (*CXCL1*, *CXCL2, Cxcl1, Cxcl2*) in primary human (A) and mouse (B) primary keratinocytes treated with BSA-conjugated free fatty acids (FFAs) ranging from C14:0 to C18:0. Untreated controls refer to cells treated with unconjugated BSA. **(C)** Gene expression of inflammatory cytokines (*Cxcl2, Ccr2*) in bone marrow derived macrophages (BMDMs) following BSA-conjugated FFAs stimulation in vitro. Untreated controls refer to cells treated with unconjugated BSA. **(D)** Representative histological sections of mouse ears treated with various FAEEs. EtOH (ethanol) treatment was used as control. Scale bars, 50 μm. **(E)** Ear thickness measurements in mice two days after topical application of FAEEs ranging from C14:0-C18:0. **(F)** Quantification of follicle size following treatment in the histology sections. **(G)** Quantification of inflammatory cell infiltration around hair follicles in the histology sections. **(H)** Ear swelling in mice topical application of C16:0 or EtOH was measured daily before treatment. **(I)** Dermoscopic images showing enlarged follicles and capillary dilation in C16:0 treated skin (left). Scale bars, 1 mm. Representative violin plots show the distribution of ring - core intensity difference (Δ) values of sebaceous glands in ear skin from EtOH and C16:0 treatment groups (right). Δ was calculated as the difference between the mean inverted grayscale intensity of the bright peripheral ring (ring_mean) and the central dark core (core_mean) of each sebaceous gland. Larger Δ values indicate stronger contrast between the glandular lumen and the surrounding rim. Data are shown as distributions with mean values indicated. Mann-Whitney U test, ∗ *p < 0.05*. **(J)** Gene expression analysis of keratinization markers (*Krt16*, *Krt17*) and inflammatory mediators (*Cxcl2*, *S100a8*, *S100a9*) in C16:0- and C18:0-treated mouse skin. For in vivo experiments (D-I), controls refer to vehicle-treated skin (EtOH, ethanol). Data points in (A)-(C) represent biological replicates and in (E)-(G) and (J) represent individual mice and bars represent mean ± SEM. All data were collected from at least two independent experiments. ∗*p* < 0.05; ∗∗*p* < 0.01; ∗∗∗*p* < 0.001; ns, not significant. Data in (A-C, E-G, J) were analyzed by one-way ANOVA with Dunnett’s T3 multiple comparisons test, and data in (H, I) by Mann-Whitney test.

To determine how long-chain saturated fatty acid (lcSFA) chain length influences classical acne pathology, we developed a mouse model of acne-like skin inflammation using topical application of fatty acid ethyl esters (FAEEs) ranging from C12 to C18. FAEEs were used to improve solubility and skin permeability and were expected to be rapidly hydrolyzed by cutaneous esterases to yield the corresponding lcSFAs, which represent the bioactive forms implicated in inflammation and comedogenesis.^12,13^ Two days after topical application of FAEEs (C14:0 to C18:0, C16:1 and C18:1) to the mouse ears, histological analysis of epidermal thickness at follicular openings revealed a chain length-dependent increase, with saturated FAEEs of C14:0 and longer, inducing significant epidermal thickening in the upper portion of the hair follicle compared to shorter-chain FAEEs and vehicle controls (**Figure 2D and 2E**). Histological assessment further showed that hair follicle size, a parameter reflecting follicular hyperkeratosis and sebum accumulation/sebocyte hyperplasia, was significantly enlarged, especially in C16:0 treated skin, (**Figure 2D and 2F**). In contrast, unsaturated C16:1 and C18:1 failed to induce such changes (**Figure 2D, S2A and S2B**). Importantly, only C16:0 triggered robust neutrophil-rich follicular inflammation, whereas C14:0 and C18:0 caused only mild responses (**Figures 2D and 2G**). Consistently, the C16:0 treated group exhibited significant overall ear thickening **(Figure 2H**). Dermoscopic image analysis revealed enhanced contrast between the perifollicular ring (Ring) and the follicular core (Core) in C16:0 treated mice, a change that may reflect enlarged follicles compared with vehicle-treated skin (**Figure 2I**).

Gene expression analysis by quantitative PCR revealed that keratinization markers *Krt16* and *Krt17* were strongly upregulated by C16:0 and C18:0 than control (**Figure 2J**). Consistent with histopathology, neutrophil-associated alarmins *S100a8* and *S100a9* were significantly induced by both C16:0 and C18:0, with the highest expression levels observed in the C16:0 treated group. Notably, the proinflammatory chemokine *Cxcl2* was elevated only in C16:0 treated skin compared with control, highlighting a more pronounced inflammatory cell related chemokine response to C16:0 than C18:0 (**Figure 2J**). In contrast, consistent with the histopathological findings, C16:1 and C18:1 induced little to no increase in gene expression (**Figure S2C**). Together with our human data, these results demonstrate that C16:0 uniquely combines comedogenesis, sebocyte hyperplasia, and neutrophil-rich inflammation, establishing it as the principal pathogenic fatty acid driving inflammatory acne.

### C16:0 exhibits higher transdermal permeability than C18:0 and promotes inflammatory cell recruitment

To directly compare the inflammatory potential of C16:0 and C18:0, we examined immune cell infiltration in mouse skin. Flow cytometry revealed a dramatic increase of Ly6C^mid^Ly6G^high^ neutrophils in C16:0 treated skin (21.24% ± 1.98%) compared with controls (3.14% ± 0.607%), whereas C18:0 induced only a modest rise (8.22% ± 1.18%) (**Figure 3A**). Similar trends were observed in the absolute cell numbers (**Figure 3A**). Ly6C^high^Ly6G^neg^ monocytes were also significantly elevated in the C16:0 group, exceeding both control and C18:0 groups (**Figure 3A**). These findings identify C16:0 as a distinct FFA that not only promotes comedogenesis but also drives neutrophil-rich follicular inflammation, closely mirroring the immune profile of human acne.

**Figure 3.**
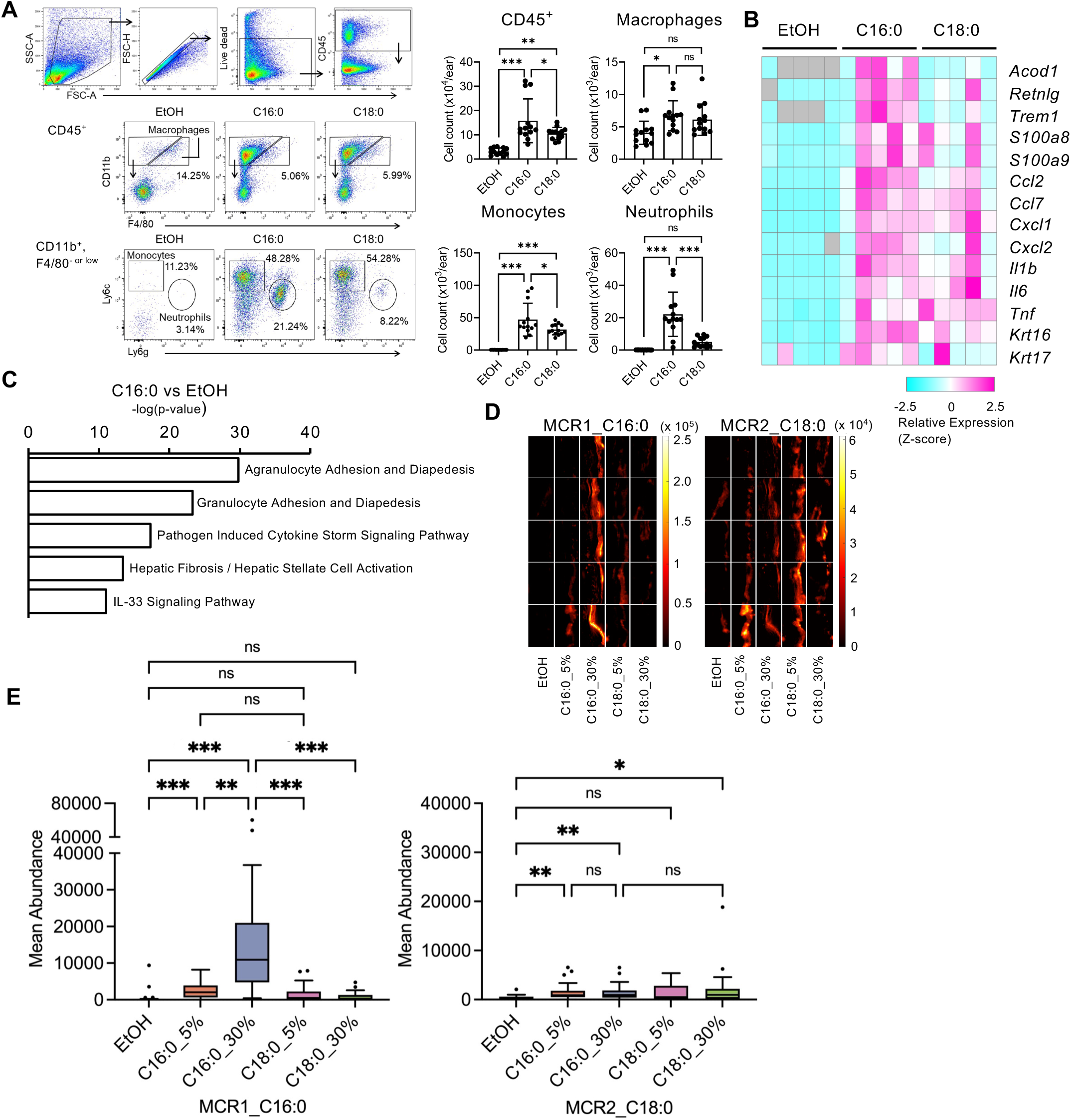
C16:0 exhibits higher transdermal permeability than C18:0 and promotes inflammatory cell recruitment. **(A)** Flow cytometric analysis of immune cell populations in skin two days after topical application of C16:0 or C18:0. The percentage of Ly6C^mid^Ly6G^high^ neutrophils (%NT) and Ly6C^+^Ly6G^neg^ monocytes (%MONO) are shown. **(B)** Bulk RNA sequencing of skin tissue collected 6 hours after topical treatment with C16:0 or C18:0, showing upregulation of genes related to keratinization (*Krt16*, *Krt17*), chemokine signaling (*Cxcl1*, *Cxcl2*, *Ccl2*, *Ccl7*), antimicrobial responses (*S100a8*, *S100a9*), and myeloid cell activation (*Acod1*, *Retnlg*, *Trem1*) in C16:0-treated skin. **(C)** Ingenuity pathway analysis (IPA) showing significant activation of pathways related to agranulocyte and granulocyte adhesion and diapedesis in C16:0-treated skin compared to untreated and C18:0-treated skin, consistent with neutrophil-dominated inflammation observed histologically and by flow cytometry. **(D)** Raman imaging and multivariate curve resolution analysis constrained by reference spectra for C16:0 (MCR1) and C18:0 (MCR2), measured on skin samples 5 minutes after topical application of C16:0 or C18:0. Images are greyscale but the “hot” colormap is used to enhance visibility of low components. 5 representative images are shown for each group from a total of 30 per class. Each small image panel represents 59 by 36 micrometers at the sample. **(E)** The reference spectra for C16:0 shows selectivity for C16:0 (MCR1 images), whereas the reference spectra for C18:0 does not. Analysis of abundances displayed in the box-whisker plots show distinctly higher abundance for C16:0 than for C18:0. Controls refer to vehicle-treated skin (EtOH, ethanol). Data points in (A) represent represent individual mice and bars represent mean ± SEM. Data in (A) were collected from at least two independent experiments. ∗*p* < 0.05; ∗∗*p* < 0.01; ∗∗∗*p* < 0.001, ns, not significant. Data were analyzed by one-way ANOVA with Dunnett’s T3 multiple comparison test. Data in (D) were collected from two independent experiments.

To investigate early transcriptional changes, we performed bulk RNA sequencing 6 hours post-treatment. In C16:0 treated skin, genes involved in keratinization (e.g., *Krt16, Krt17*), chemokine signaling (e.g., *Cxcl1, Cxcl2, Ccl2, Ccl7*), neutrophil-associated alarmins (e.g., *S100a8, S100a9*) and myeloid cell activation (e.g., *Acod1, Retnlg, Trem1*) were significantly upregulated compared with vehicle-treated controls, with the increase more pronounced in contrast to the C18:0-treated group (**Figure 3B**). Ingenuity pathway analysis (IPA) further revealed that the most significantly enriched pathways after C16:0 treatment included both agranulocyte and granulocyte adhesion and diapedesis, supporting the histological and flow cytometric findings of neutrophil/monocyte-dominated inflammation (**Figure 3C**).

Interestingly, our in vitro assays in this study suggest stronger activity for C18:0 in keratinocytes and BMDMs. To resolve this discrepancy, we quantified lipid penetration using Raman imaging and multivariate curve resolution (MCR) analysis, constrained to either C16:0 (MCR1) or C18:0 (MCR2) reference spectra. FFAs were applied topically at 5% or 30%, and Raman imaging was performed after 5 minutes (**Figure 3D**). In MCR1, both 5% and 30% C16:0 showed markedly higher signals than controls, with 30% exceeding 5% (*p* = 0.0065). Both concentrations were significantly greater than any C18:0 condition (all *p* < 0.0001). In contrast, MCR2 revealed only weak signals for C18:0; although 30% C18:0 showed a modest increase vs. control, it did not differ from C16:0 at either concentration (**Figure 3E**).

Together, these analyses demonstrate that C16:0 penetrates skin tissue far more effectively than C18:0, with greater signal intensity and concentration-dependent enhancement when detected against its own spectral signature, whereas C18:0 shows only minimal enrichment even under high-concentration conditions.

### C16:0 activated *Lrig1^+^* cells drive inflammation and hyperkeratinization within a primed follicular niche

To further characterize the cellular sources of these inflammatory signals, we performed single-cell RNA sequencing (scRNA-seq) of skin from control and C16:0-treated mice, collected at 6 and 12 hours post-treatment (**Figure S3A**). Through UMAP visualization, unsupervised clustering identified 14 distinct cell populations (clusters 0-13) encompassing major skin-resident cell types (**Figure S3B and S3C**). Within the myeloid compartment (cluster 3), comprising of macrophages, monocytes, neutrophils, and NK cells, all populations expanded following C16:0 treatment, with monocytes and neutrophils showing the most pronounced increases (**Figure S3D and S3E**), consistent with histological and flow cytometry findings.

We next focused on epithelial populations to identify the sources of chemokine expression driving immune cell recruitment by analyzing *Cxcl* and *Ccl* expressions across epithelial populations by scRNA-seq. At baseline, the *Lrig1^+^Scd1^+^Mgst1^+^* sebocyte cluster (SEBII population in cluster 0, **Figure 4A and 4B**) constitutively expressed *Cxcl1* (**Figure 4C**). Upon C16:0 treatment, *Cxcl1, Cxcl2, Ccl2,* and *Ccl20* expressions were further upregulated in these *Lrig1^+^* sebocytes (**Figure 4C**). In addition, *Lrig1^+^Krt79^+^* follicular keratinocytes (HF-KCII population in cluster 10, **Figure 4A and 4B**) also expressed *Cxcl1* and *Ccl20* following C16:0 application (**Figure 4C**). In contrast, epidermal keratinocytes (KC population, **Figure 4A and 4B**) showed only weak *Ccl20* expression following C16:0 stimulation (**Figure 4C**). These findings provide a mechanistic explanation for why inflammatory cell infiltration occurs selectively in hair follicles rather than diffusely across the epidermis in response to C16:0.

**Figure 4.**
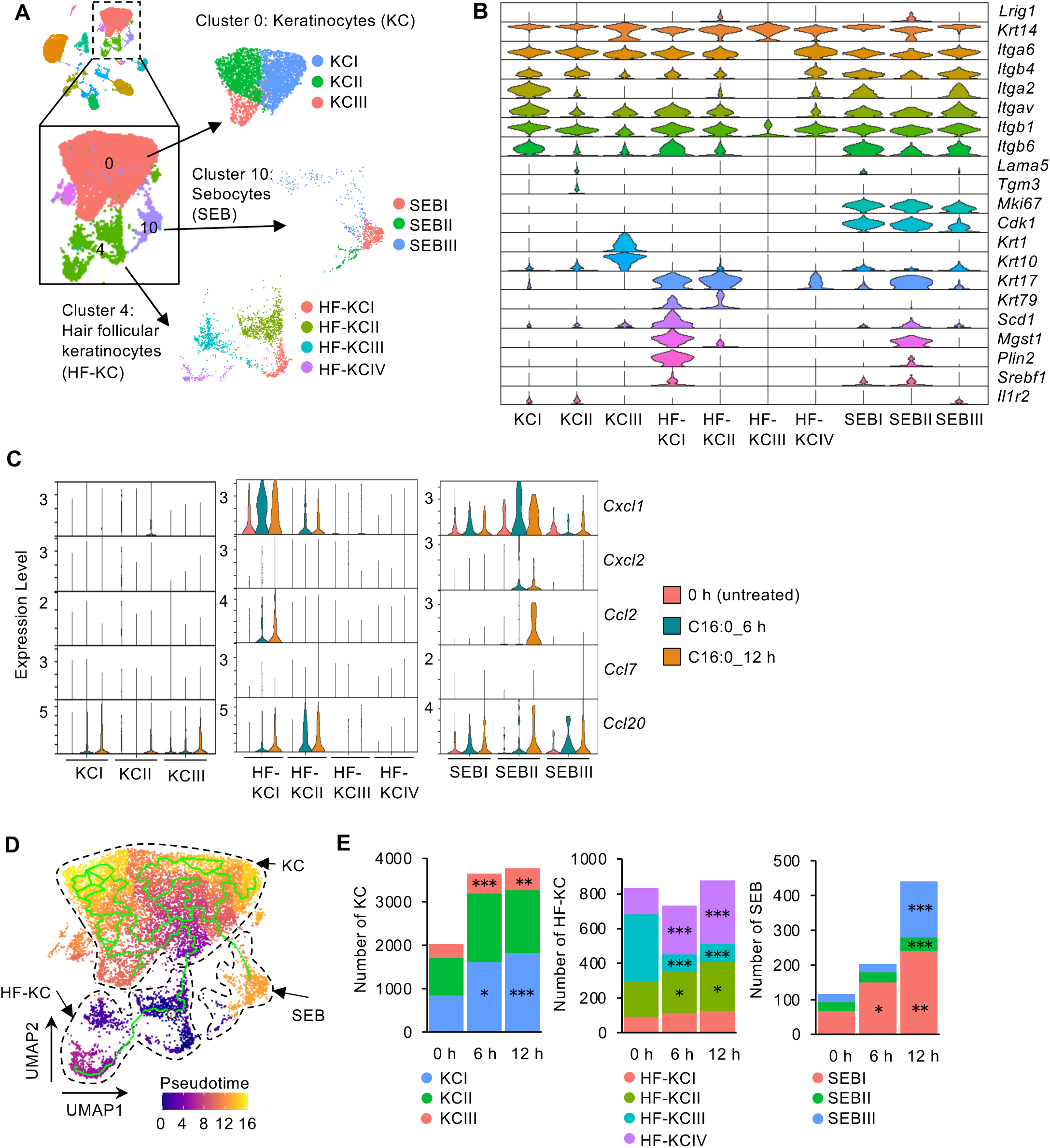
C16:0-activated *Lrig1* cells drive inflammation and hyperkeratinization within a primed follicular niche. **(A)** Overview of scRNA-seq clustering and subpopulation analysis. A UMAP of all 14 skin cell clusters (shown in Figure S3C) is displayed in the upper left corner. The dotted box highlights the epithelial populations, keratinocytes (KC, cluster 0), hair follicular keratinocytes (HF-KC, cluster 4), and sebocytes (SEB, cluster 10). These populations are shown enlarged in the main panel and further subdivided into color-coded subpopulations (KCI-III, HF-KCI-IV, SEBI-III), which are displayed as separate plots at the arrowheads. **(B)** Expression levels (y axis) of major cluster-defining genes in each cluster. Violin plots show the distribution of the normalized expression levels of genes. **(C)** Chemokine expression profiles (*Cxcl1, Cxcl2, Ccl2*, and *Ccl20*) after C16:0 treatment in keratinocytes (cluster 0), sebocytes (cluster 10) and hair follicular keratinocytes (cluster 4). **(D)** Pseudotime analysis showing the differentiation trajectory originating from *Lrig1^+^* progenitor cells (HF-KCII and SEBII), giving rise to *Lrig1* sebocytes (cluster 10), hair follicular keratinocytes (cluster 4, KCI-III), and keratinocytes (cluster 0). **(E)** Quantification of epitherial cell subsets sebocytes (cluster 10), hair follicular keratinocytes (cluster 4, HF-KCI-IV), and keratinocytes (cluster 0, KCI-III) derived from scRNA-seq data. Data are from a single experiment with five biological replicates (individual mice). Statistical significance in (E) was assessed by chi-square test followed by analysis of standardized residuals to identify cell populations contributing to overall distribution changes. ∗*p* < 0.05; ∗∗*p* < 0.01; ∗∗∗*p* < 0.001.

To explore lineage dynamics, pseudotime analysis revealed a differentiation trajectory originating from *Lrig1^+^* progenitor cells, which gave rise to *Lrig1^−^* cluster 10 sebocytes and cluster 0 keratinocytes after C16:0 application (**Figure 4D and 4E**). Notably, in the sebocyte (SEB) population, the number of cells increased approximately four-fold within 12 hours after stimulation, indicating a marked expansion of differentiated sebocytes (**Figure 4E**). This cellular increase was consistent with histopathological observations showing sebaceous hyperplasia after C16:0 stimulation. These findings suggest that C16:0-stimulated *Lrig^+^* progenitor cells not only initiate localized inflammation but, by supplying differentiated sebocytes, also establish a primed follicular niche capable of amplifying inflammatory signaling by supplying differentiated sebocytes.

### C16:0 induced inflammatory comedones are independent of microbiota and fatty acid transporter function

To elucidate the mechanisms underlying the activation of *Lrig1^+^*progenitor cells by C16:0, we performed Kyoto Encyclopedia of Genes and Genomes (KEGG) pathway enrichment analysis on the differentially expressed genes within this population. The analysis revealed significant enrichment of multiple signaling pathways, including endocytosis, MAPK, PI3K, and Ras pathways, suggesting that C16:0 is involved in diverse intracellular signaling cascades in these cells (**Figure 5A and 5B**). In addition, enrichment of lipid metabolic pathways was observed, most notably the sphingolipid signaling pathway, which includes ceramide biosynthesis and showed a highly significant enrichment (adjusted p value = 5.56 × 10 ¹) (**Figure 5B**).

**Figure 5.**
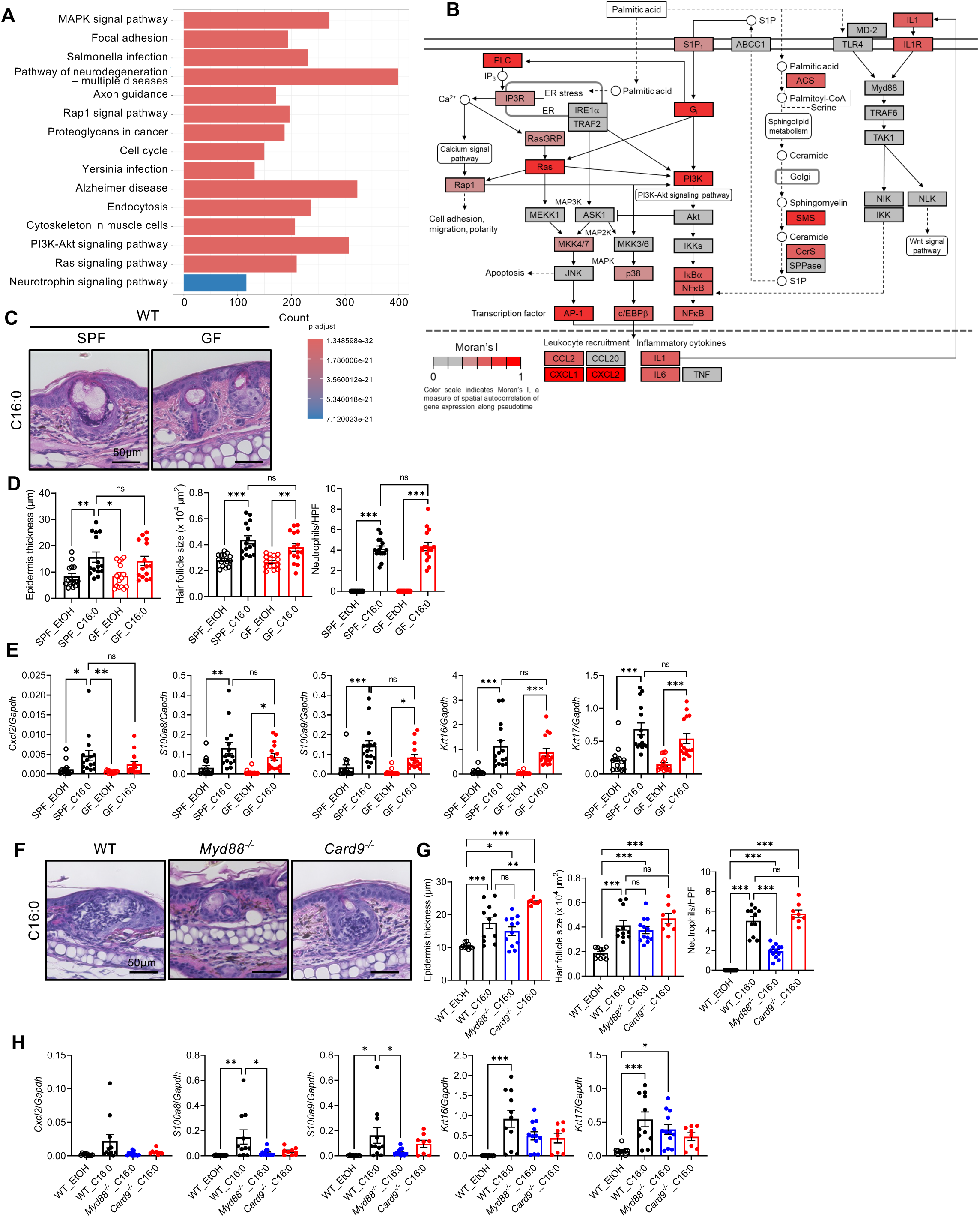
C16:0 induced inflammation is microbiota-independent and partially Myd88-dependent. **(A)** KEGG pathway analysis of *Lrig1* progenitor cells from scRNA-seq data showing the top 15 significantly enriched pathways following C16:0 treatment. **(B)** Schematic representation of KEGG pathways relevant to *Lrig1* progenitor cell responses to C16:0, including ER stress, MAPK, PI3K, AP-1, Ras, Sphingolipid signaling and TLR/IL1R pathways. The schema was modified from KEGG pathway maps (KEGG, https://www.genome.jp/kegg/). **(C)** Representative histological sections of C16:0 treated skin from wild-type (WT) mice maintained under germ-free (GF) or specific pathogen-free (SPF) conditions. **(D)** Measurements of ear thickness, follicle size, and inflammatory cell infiltration in GF and SPF mice following topical C16:0 application. **(E)** Gene expression of *Krt16, Krt17, Cxcl2, S100a8* and *S100a9* in C16:0 treated skin from GF and SPF mice. **(F)** Representative histological sections of ear thickness measurements in wild-type (WT), *Myd88*-deficient (*Myd88^-/-^*), and *Card9*-deficient (*Card9^-/-^*) mice following C16:0 application. **(G)** Measurements of ear thickness, follicle size, and inflammatory cell infiltration in *Myd88^-/-^* and *Card9^-/-^* mice following C16:0 application. **(H)** Gene expression of *Krt16, Krt17, Cxcl2, S100a8* and *S100a9* in C16:0 treated skin from WT, *Myd88^-/-^,* and *Card9^-/-^* mice. Data in (A and B) are from a single experiment with five biological replicates (individual mice). Data points in (D-F, H-J) represent individual mice, and bars indicate mean ± SEM. Data were collected from at least two independent experiments. Controls refer to vehicle-treated skin (EtOH, ethanol). *p* < 0.05; *p* < 0.01; *p* < 0.001; ns, not significant. statistical analysis by one-way ANOVA with Dunnett’s T3 multiple comparisons test.

To evaluate the contribution of the skin microbiota to C16:0-induced inflammatory comedones, we applied C16:0 topically to the ears of germ-free (GF) mice. Despite the absence of commensal microbes, GF mice exhibited comparable ear thickening to that observed in specific pathogen-free (SPF) mice (**Figure 5C and 5D**). Histological analysis further revealed that C16:0-treated GF mice developed follicular hyperkeratosis and sebocyte hyperplasia/sebum accumulation like its SPF counterparts. Gene expression levels of *Cxcl2, S100a8, S100a9, Krt16*, and *Krt17* were also comparable between the two groups **(Figure 5E)**, suggesting that C16:0 driven folliculitis occurs largely independent of the presence of cutaneous or gut microbiota.

Given prior reports that lcSFAs such as C16:0 can act as endogenous ligands for the TLR4–MD2 complex,^14,15^ we next examined the role of downstream signaling. *Myd88*-deficient mice, as well as *Card9*-deficient mice (defective in C-type lectin receptor signaling), were used to define potential innate immune pathways. Consistent with this approach, neither *Myd88-* nor *Card9-*deficient mice, lacking key adaptor proteins involved in microbial recognition, exhibited reductions in C16:0 induced sebocyte hyperplasia or epidermal hyperkeratosis compared to wild-type controls (**Figure 5F and 5G**). In contrast, inflammatory cell infiltration was markedly suppressed in *Myd88-*deficient mice, indicating a specific role of *Myd88* signaling in C16:0-triggered inflammation. Supporting this, gene expression analysis revealed that while *Krt16* and *Krt17* levels remained comparable to wild-type in both *Myd88-* and *Card9-*deficient mice, *Cxcl2* and *S100a8/a9* expressions were modestly reduced in *Myd88-*deficient skin (**Figure 5H**). However, further analysis using keratinocyte-specific (*K14CreMyd88^f/f^*) and myeloid-specific (*LysMCreMyd88^f/f^*) conditional knockout mice showed no significant differences in ear histopathology, or gene expressions compared to wild-type mice (**Figures S4A, S4B and S4C**), consistent with a requirement for Myd88 signaling across multiple cell types. These results demonstrate that C16:0-induced folliculitis is independent of both cutaneous and gut microbial signals and is driven primarily by the FFA itself. While sebocyte hyperplasia and keratinization appeared to be TLR4/MyD88-independent, the priming and recruitment of inflammatory cells partially required TLR4/Myd88-dependent signaling.

We next investigated whether C16:0-induced inflammation depends on fatty acid transporter function. Using our scRNA-seq dataset, we assessed the expression of known fatty acid transporters in *Lrig1^+^* progenitor cells, the primary target population of C16:0. Among these, only *Fabp5* and *Slc27a3* demonstrated detectable expression, whereas other transporters (*Fabp3, Fabp4, Cd36, Slc27a1, Slc27a4, Slc27a6*) did not (**Figure S5A**). To examine the functional relevance of Fabp5, we analyzed C16:0 induced responses in Fabp5-deficient mice and observed no significant differences in histology or inflammatory gene expression compared to wild-type controls (**Figures S5B, S5C and S5D**). Slc27a3, also known as fatty acid transport protein 3 (Fatp3), has been reported to be activated by 3-hydroxy-isobutyrate (3-HIB).^16^ However, oral administration of 3-HIB did not exacerbate C16:0 induced folliculitis, with no detectable changes in tissue swelling, histopathology, or gene expression (**Figures S5E-G**).

In parallel, scRNA-seq analysis of *Lrig1^+^* progenitor cells following C16:0 treatment revealed enrichment of stress-related pathways, including JNK signaling, endocytosis, and ER stress **(Figure 5A, 5B)**. Together, these findings suggest that the inflammatory effects of C16:0 in this model are not dependent on fatty acid transporter activity, but are likely mediated through multiple mechanisms, potentially involving the TLR/MyD88 axis as well as endocytosis- and stress-associated signaling pathways.

## Discussion

In this study, we identified C16:0 (palmitic acid), the most abundant lcSFA in human skin, as a key pathogenic lipid that drives initial comedogenesis and subsequent follicular inflammation. Using human lipid profiling, a novel mouse model, and single-cell transcriptomics, we demonstrated that C16:0 selectively accumulates in acne-prone skin, penetrates the epidermis, and triggers inflammatory and differentiation programs within *Lrig1^+^* hair follicle progenitor cells. These responses occurred independently of microbial cues or fatty acid transporter activity, revealing a direct lipid-driven mechanism of skin inflammation.

A clear structure–activity relationship was observed: comedogenic potential increased with carbon chain length, but only C16:0 consistently reproduced acne-like folliculitis features in vivo. However, these effects are unlikely to depend on a single receptor. Instead, C16:0 has been shown in non-cutaneous systems to activate multiple converging mechanisms amplifying inflammation. These include altered membrane fluidity and lipid raft organization, which facilitate TLR4 clustering and downstream signaling,^17,18^ induction of ER stress and JNK/AP-1 activation,^19,20^ and ceramide biosynthesis–driven inflammatory signaling.^21–23^

Despite skin being one of the most lipid-rich organs, a systematic model to explore lipid-driven inflammation has yet to be established. In this study, by a topical application of C16:0, we established a mouse model that closely recapitulates key features of human acne, including sebocyte hyperplasia, comedogenesis, and neutrophil-rich follicular inflammation, in a manner consistent with the proinflammatory properties of C16:0 observed in other tissues.

Raman spectroscopy further demonstrated that C16:0 efficiently penetrated the skin, whereas C18:0 did not. Previous studies have shown that Raman spectroscopy can visualize and quantify lcFFAs in cells and is widely used to analyze cellular components, including fatty acids.^16,24^ Nonetheless, most tissue-based applications mainly focused on broader classifications, such as distinguishing lipids from proteins. A few exceptions include attempts to quantify drug penetration^25^ and depth profiling in ex vivo tissue using water content signatures in the high-wavenumber region.^26,27^ In this study, we leveraged Raman imaging across 150 tissue samples, treated with varying concentrations of FFAs and using measured reference spectra to quantify the relative abundance of each FFA within the tissue. These findings highlight that in vivo bioactivity depends not only on intrinsic inflammatory potential but also on cutaneous bioavailability.

One of the longstanding questions in the pathogenesis of acne is whether inflammation is primarily initiated by host-derived FFAs or microbial metabolites. In our study, we found that odd-chain FFAs such as C15:0 and C17:0, FFAs predominantly produced by skin commensals, were present at only trace levels in both acne-prone and healthy human skin. Moreover, in a mouse model, C16:0 induced inflammation was comparable between GF and SPF conditions, suggesting that microbial colonization is not essential for triggering the inflammatory response. These findings indicate that host-derived lcSFAs, rather than microbial products, are the primary drivers of inflammation and comedogenesis. This supports the emerging concept that acne is fundamentally an autoinflammatory disorder mediated by lipid dysregulation, rather than a disease caused by microbial overgrowth. Importantly, our findings are consistent with the ongoing clinical shift away from antibiotic-based acne treatments toward strategies that target keratinocyte turnover and inflammatory signaling pathways.^28^ Furthermore, they may provide mechanistic insight into why minocycline is particularly effective in some patient subsets, as it exhibits anti-neutrophilic properties, such as inhibiting neutrophil chemotaxis and suppressing matrix metalloproteinase activity, independent of its antimicrobial action.^29^

Notably, C16:0-induced inflammation was preserved in germ-free mice and in *Myd88* deficient and *Card9* deficient mice, including keratinocyte- and myeloid-specific *Myd88* deficient mice. This microbiota-independent response challenges the prevailing view that lcSFAs act primarily through TLR4 signaling. Although Myd88 deficiency led to a modest reduction in inflammatory gene expression and a mild decrease in tissue inflammation, the overall phenotype remained largely intact, suggesting that TLR4 signaling may play only a secondary role. This is consistent with recent findings that palmitate does not directly activate TLR4, but instead induces ER stress, which may secondarily amplify inflammatory pathways.^30^ Importantly, candidate lipid transporters expressed in *Lrig1* progenitor cells, such as Fabp5 and Slc27a3 (Fatp3), were tested using knockout mice or ligand-based modulation. In both cases, C16:0 driven comedogenesis and inflammation persisted, arguing against a critical role for these transporters. Taken together, these findings indicate that the pathogenic activity of C16:0 is not mediated by a single receptor or transporter but instead emerges from multiple converging stress- and signaling pathways.

Consistent with these observations, scRNA-seq revealed that C16:0 activates ER stress, JNK signaling and ceramide biosynthesis in *Lrig1^+^* progenitor cells, supporting a stress-mediated, microbe-independent mechanism of inflammation. Furthermore, *Lrig1^+^*progenitor cells have previously been identified as quiescent stem-like populations capable of multilineage differentiation,^31^ and our findings further extend their functional role to lipid sensing and immune modulation. These cells expanded and differentiated into both sebocyte and keratinocyte lineages, consistent with sebaceous hyperplasia and follicular hyperkeratinization. They also expressed high levels of chemokines such as *Cxcl1, Cxcl2*, and *Ccl20*, which drive neutrophil/monocyte recruitment. In contrast, interfollicular epidermal keratinocytes lacked this chemokine response, offering a mechanistic explanation for the spatial confinement of inflammation to the pilosebaceous unit. Togethre, these results suggest that *Lrig1^+^* progenitor cells function not only as structural stem-like cells but also as dynamic sensors of lipid cues, capable of translating metabolic stimuli into immune activation within the pilosebaceous niche.

These findings support a revised model of acne pathogenesis in which endogenous saturated lipids, rather than microbial factors, serve as primary initiators of disease, opening new therapeutic avenues. Targeting C16:0 biosynthesis, uptake, or downstream stress-response pathways may represent promising strategies. Furthermore, *Lrig1^+^* progenitor cells constitute a previously unrecognized therapeutic target in acne, functioning as both lipid sensors and amplifiers of inflammation. Future studies should therefore explore whether modulation of this niche can attenuate comedogenesis and restore follicular homeostasis.

Overall, this work establishes a mechanistic link between skin lipid composition and localized inflammatory pathology, providing a foundation for lipid-centered strategies in the diagnosis and treatment of acne and potentially other inflammatory skin diseases.

## MATERIALS and METHODS

### Contact for Reagent and Resource Sharing

Further information and requests for resources and reagents should be directed to and will be fulfilled by the corresponding author, Yuumi Nakamura. Any additional information required to reanalyze the data reported in this work paper is available from the lead contact upon request.

### Experimental model and study participant details

#### Clinical study design

A total of 43 subjects with severe-to-mild acne vulgaris (female, age range 20–39, mean 27.4) and seven healthy subjects (female, age range 21–37, mean 32.1) were included in the study. We obtained human skin sebum samples from the forehead skin of acne vulgaris and healthy subjects. The study protocols were approved by The University of Osaka (Osaka, Japan, Approval Numbers: 20390, 21071) and Rohto Pharmaceutical Co., Ltd. Clinical Research Ethics Review Committee (Osaka, Japan, Approval Numbers: ROHTO-20-024, ROHTO-21-020, ROHTO-21-0203). Informed consent was obtained from all participants involved in the study. The classification of healthy individuals and acne vulgaris patients was conducted by The Japanese Dermatological Association board-certified dermatologists. The numbers of inflammatory and non-inflammatory lesions were also evaluated by the dermatologists. Disease severity was determined based on the number of inflammatory lesions, in accordance with the Japanese Dermatological Association guidelines for acne vulgaris.

#### Animals

C57BL/6J, 6N and BALB/cAJcl mice were purchased from CLEA Japan (Tokyo, Japan). *K14-Cre* mice were purchased from the Jackson Laboratory. *LysM-Cre* mice were provided by the RIKEN BRC through the National BioResource Project of the MEXT/AMED, Japan. *Myd88^-/-^*, *Card9^-/-^*mice (C57BL/6 background) have been described previously,^32,33^ and were bred and maintained in the animal facilities of the University of Osaka. *Myd88^fl/fl^* mice (C57BL/6 background) were kindly provided by Dr. Gabriel Nunez (The University of Michigan). *Fabp5^-/-^* mice (C57BL/6 background) were kindly provided by Dr. Motoko Maekawa (Tohoku University) *Myd88^fl/fl^* mice were crossed with *K14-Cre* mice or *LysM-Cre* mice to generate *K14CreMyd88^fl/fl^* (keratinocyte-specific knockout) mice or *LysMCreMyd88^fl/fl^* (myeloid-specific knockout) mice. All above mice were bred and/or maintained under the specific pathogen-free facility at The University of Osaka in accordance with the Guide for the Care and Use of Laboratory Animals. Germ-free C57BL/6NJcl mice were purchased from CLEA Japan. Female aged-matched mice between 7 and 12 weeks of age were used for each experiment. All animal experiments were performed in strict accordance with the regulation of Animal Experimentation at The University of Osaka. The protocols (No.03-028-017, 03-028-020) were approved by The University of Osaka Institutional Animal Care and Use Committee.

#### Cells and cell culture

Normal human epidermal keratinocytes (NHEKs) were obtained from Kurabo (Osaka, Japan) and maintained in HuMedia-KG2 culture medium (Kurabo) supplemented with 5 μg/ml insulin, 0.5 μM hydrocortisone, 200 pg/ml human recombinant epidermal growth factor, and 0.2% bovine pituitary extract (growth medium). Murine primary keratinocytes were isolated from neonatal C57BL/6 J mice (postnatal day 1-3). Skin tissues were incubated with 5 mg/mL dispase (CELLnTEC) at 37 °C for 30 minutes to separate the epidermis from the dermis. The isolated epidermal sheets were then treated with accutase (CELLnTEC) at room temperature for 20 minutes. Single-cell suspensions were prepared and cultured in CnT-Prime epithelial culture medium (CELLnTEC) for 5-6 days. Bone-marrow-derived macrophages (BMDMs) were isolated from C57BL/6 J mice and maintained using Iscove’s Modified Dulbecco’s Medium (IMDM, Sigma) supplemented with 10% fetal bovine serum (FBS, Sigma), 30% L-cell conditioned medium, 1% MEM non-essential amino acids, 1% sodium pyruvate, 0.025 mM 2-mercaptoethanol and 1% penicillin-streptomycin. All cells were maintained at 37°C in a humidified atmosphere at 5% CO2.

### METHOD DETAILS

#### Sebum free fatty acid analysis

Sebum samples were collected from the center of the forehead area of both healthy individuals and acne vulgaris patients using Sebutape^®^ (CuDerm Corp), which was applied for 30 minutes. Sebutape^®^ is a microporous film designed for the collection of sebum from the skin. FFAs collected on the Sebutape® were subsequently extracted by using the Folch method. ^34^ In brief, the tape was detached from the backing paper and placed in a glass test tube containing 8 ml of extraction solvent (2:1 mixture of chloroform and methanol) along with glass beads. The mixture was shaken for 10 minutes, followed by 10 minutes of sonication, and then shaken for an additional 10 minutes to extract FFAs from the sebum. An unused Sebutape^®^ served as the blank control. The solvent containing the extracted FFAs was evaporated under a nitrogen gas stream, and the residue was dissolved in 1 ml of a chloroform solution containing 20% ethanol (w/v). Subsequently, 0.5 ml of trimethylsilyldiazomethane (approximately 0.6 mol/L hexane, Tokyo Chemical Industry Co., Ltd.) was added, thoroughly mixed, and allowed to react at room temperature for 30 minutes. The solvent from the resulting solution was evaporated under a nitrogen gas stream. An internal standard (C24 ester) was introduced, and the solution was transferred to a glass vial where the solvent was evaporated under a nitrogen gas stream. The residue was dissolved in 200 μl of hexane and analyzed using Gas Chromatography/Mass Spectrometry.

#### Preparation of BSA-conjugated fatty acids

Preparation of BSA-conjugated fatty acids was performed as previously described^35^, with minor modifications. Fatty acids (all purchased from Sigma): lauric acid (C12:0), myristic acid (C14:0), palmitic acid (C16:0), and stearic acid (C18:0) were dissolved in 100% ethanol to prepare 250 mM stock solutions by vortexing until fully solubilized. A 10% (w/v) fatty acid-free bovine serum albumin (BSA) solution was prepared by dissolving fatty acid -free BSA (Sigma) in distilled water, followed by mixed gently by vortexing and incubation at 37 °C for at least 15 minutes. The pH was adjusted to 7.4-7.5 using NaOH, and the solution was sterilized through a 0.22 μm filter. To generate the fatty acid-BSA complex, the 250 mM fatty acid stock was added to the 10% BSA solution to achieve a final fatty acid concentration of 9 mM. During addition, the mixture was gently stirred with a vortex mixer to prevent aggregation and precipitation. The final mixture was incubated at 37 °C for 30 minutes with intermittent vortexing to ensure thorough solubilization and complex formation.

#### Stimulation of cultured cells with BSA-conjugated fatty acids

NHEKs, murine primary keratinocytes, and BMDMs were seeded into 24-well plates (growth area: 2 cm²/well). Seeding densities were as follows: NHEKs at 5.0 × 10 cells/cm², murine primary keratinocytes at 4.0 × 10 cells/cm², and BMDMs at 2.5 × 10 cells/cm². BSA-conjugated fatty acids were added to the culture medium at a final concentration of 500 μM. As a control, BSA alone (without fatty acids) was used. After 6 hours of treatment, cells were collected for gene expression analysis.

#### Mouse model of skin folliculitis by fatty acids

Each FAEE was prepared as a 5% (w/w) solution in ethanol. For the topical application model, 20 µL of each solution was applied to the mouse ears every 24 hours for two consecutive days.

#### Image preprocessing and feature extraction

Dermoscopic images of the mouse ear skin were acquired using a dermocamera (DZ-D100, Casio) with a fixed magnification of 7.36x under standardized illumination. Images were contrast-enhanced using the Contrast Limited Adaptive Histogram Equalization (CLAHE) algorithm implemented in OpenCV (v4.8, Open Source Computer Vision Library). Sebaceous glands were detected as circular dark blobs with bright rims using the SimpleBlobDetector function in OpenCV, optimized for diameters of 20-60 pixels.

#### Gene expression analysis

Total RNAs were isolated from cells and mice ears using TRIzol® Reagent (Invitrogen) and the Direct-zol RNA Miniprep Kit (Zymo Research) according to the manufacturer’s instructions. RNA concentration was measured using CytationTM3 microplate reader (BioTek). Reverse transcription of RNA extracted from mouse ears was carried out using the High-Capacity cDNA Kit (Applied Biosystems), followed by quantitative real-time PCR (qPCR) to assess genes expression. The qPCR was conducted using an ViiA7 real-time PCR system (Life Technologies Corporation). Gene expression was normalized to the expression of housekeeping gene *GAPDH* or *Gapdh* mRNA.

#### Histology and immunohistochemistry

Tissues were fixed with 10% Formalin Neutral Buffer Solution (FUJIFILM Wako Pure Chemical Corporation) and then embedded in paraffin. Sections with a thickness of 4 μm were prepared and stained with hematoxylin and eosin. The images were captured using an HS All-in-one Fluorescence Microscope BZ-X710 (Keyence). Histopathological scoring was performed based on the number of infiltrating neutrophils around individual hair follicles. Neutrophils were counted in high-power fields (HPF) with a focus on their accumulation around single follicles. A semi-quantitative score was derived from the cell counts per HPF. In addition, the epidermal thickness in the upper portion of the hair follicle and hair follicle size was quantitatively measured using ImageJ software (NIH). Hair follicle size was measured as the total size of the follicle including the associated sebaceous gland. Each data point (dot) represents the value from one mouse, calculated as the average of at least three independent measurement sites.

#### Flow cytometry analysis in mouse skin

Skin tissue processing was performed as previously described by Kobayashi et al.^36^ with minor modifications. Briefly, mouse ear skin was harvested and placed in PBS on ice. The skin tissue was minced in RPMI medium containing 125 μg/mL Liberase TM (Roche) and 50 μg/mL DNase I (Sigma) in gentleMACS™ C Tubes (Miltenyi Biotec), followed by incubation at 37 °C for 2 hours. Mechanical dissociation was performed using the gentleMACS™ Dissociator to generate single-cell suspensions. Following dissociation, cell suspensions were filtered through a sterile 100 μm cell strainer. The cells were washed with HBSS containing 0.01% BSA and were further filtered through a sterile 40 μm cell strainer into a new 5ml round bottom tube. For viability and surface staining, cells were first stained with Fixable Viability Dye eFluor™ 780 (Invitrogen) for 30 minutes at room temperature, followed by Fc receptor blocking with anti-mouse CD16/32 antibody. Subsequently, cells were stained with the following fluorochrome-conjugated antibodies for 20 minutes on ice: CD45 (I3/2.3), CD11b (M1/70), F4/80 (BM8), Ly-6G (1A8), Ly-6C (HK1.4). The antibodies were purchased from BioLegend, eBioscience or BD Biosciences. The data was acquired by BD FACSCelesta™ flow cytometer (BD Biosciences) and analyzed by using FlowJo software (FlowJo, LLC). Viability Dye-stained dead cells and doublet cells were removed from the analysis and CD45^+^ cells were gated as hematopoietic cells. CD11b^+^ cells were gated as myeloid cells and CD11b^+^F4/80^+^ cells were gated as macrophages. In CD11b^+^F4/80^-^ population, Ly-6G^+^ were gated as neutrophils and Ly-6G^-^Ly-6C^+^ were gated as monocytes.

#### Bulk RNA-sequencing analysis

Total RNAs were isolated from mice ears using TRIzol® Reagent and the Direct-zol RNA Miniprep Kit according to the manufacturer’s instructions. RNA quantity was assessed using Qubit fluorometer (Thermo Fisher Scientific), and RNA integrity was evaluated using the Agilent 2100 Bioanalyzer (Agilent Technologies). Samples having a RIN value around 7 were used for further analysis. For the construction of bulk RNA-Seq library, Poly(A) mRNA Magnetic Isolation Module (New England Biolabs) and NEBNext® Ultra™II Directional RNA Library Prep Kit for Illumina (New England Biolabs) was used according to the manufacturer’s instructions and then sequenced with NovaSeq 6000 (Illumina Inc.). FASTQ files were first processed with Trimmomatic v.0.33 (Bolger et al. 2014) for adapter removal and quality filtering. The cleaned reads were mapped to the mouse transcriptome (Mus_musculus.GRCm39.cdna.all) and then quantified using Salmon (version 1.10.2). Bulk RNA-seq analysis was performed with GENE-E (version 3.0.215), which was matrix visualization and analysis software, the DESeq2 package (version 1.30.1) and Qiagen Ingenuity Pathway Analysis (IPA) software (version 81348237; Qiagen).

#### Ingenuity pathway analysis

Upstream analysis was performed using the Ingenuity Pathway Analysis (IPA) software (version 81348237; Qiagen). Differentially expressed genes were subjected to “Core Analysis,” and mapped to the Ingenuity Knowledge Base. Molecule types including genes, RNAs, and proteins were included in the upstream regulator analysis. A right-tailed Fisher’s exact test was used to calculate p-values reflecting the statistical significance of the overlap between the input dataset and known upstream regulators. Adjusted p-values were calculated using the Benjamini– Hochberg (BH) method to correct for multiple testing.

#### Raman imaging and analysis

Tissue sections from BALB/cAJcl mice were placed on quartz bottomed cell culture dishes (FPI, Japan) and covered with D-PBS(-) (Nacalai, Japan) for Raman measurements to prevent heating. Raman images were collected using a Raman-11 microscope system (Nanophoton, Japan) at 532 nm excitation, using a x40/NA 0.8 water immersion lens (Nikon, Japan), and 600 l/mm grating (resulting in a spectral range of 524-2974 cm^-1^). Images were collected by rapidly scanning the laser over a 130 pixel area for 4 s per line, with a laser power of ∼165 mW at the sample. Images were recorded from tissue sections from two mice for each treatment type, and images were taken from both the upper and lower edges of the tissue sections. Each image was 115 x 75 pixels, equating to an image area of 58.9 x 35.5 µm

Reference spectra of ethyl palmitate and ethyl stearate were recorded in point mode, using a x20/NA 0.45 dry lens (Nikon, Japan), with 1 second exposure time and 50 mW power at the sample. 7 spectra were recorded from different areas of the sample and averaged together to produce a single reference spectrum for each fatty acid. Raman images were imported into MATLAB R2019b (Mathworks, USA), then were baseline corrected using a 4th order weighted least squares algorithm without smoothing, and cosmic rays were removed by despiking with a median filter (threshold 2 and window 3), using PLS Toolbox (version 9.2.1) and MIA (version 3.0.9) toolboxes (Eigenvector Research Inc. USA). The data was then processed in Python (v3.12), SciPy (v1.13.1), and Pandas (2.2.2). MCR-ALS was calculated using nnls with non-negative constraints, with a 5 component model (the component number having been selected empirically to prevent redundancy in emergent spectral components). The first two components MCR1 and MCR2 were constrained as the reference spectra taken from C16:0, and C18:0 respectively. The remaining three were unconstrained. After convergence, the components were normalized using the L1 norm and the scaling factors required to normalize the components were transferred to the corresponding abundances, providing an estimate for the relative amounts of different components. To quantify abundances and compare between groups, each image panel was then averaged to create a single mean abundance for that measured sample for each MCR component. The statistics on the abundance values were calculated in SciPy (v1.13.1), and mean abundance box-whisker plots were generated in Seaborn (0.13.2).

#### Single-cell RNA-sequencing analysis

Following the protocol outlined for the mouse model of skin folliculitis induced by FAEEs, we applied 5% (wt/wt) ethyl palmitate (C16:0) in ethanol to mice, while control mice remained untreated (0 hour). Cells were isolated from the ears of mice at 6- and 12-hours post-application of C16:0, as well as from untreated control mice. Tissue processing was performed using the same protocol as described in the flow cytometry section. 5 mice were pooled per group to perform sorting with the cell sorter SH800ZFP (SONY). Cells were first stained with Fixable Viability Dye eFluor™ 780 (eBioscience, Invitrogen) for 30 minutes at room temperature. Fc receptors were then blocked using anti-mouse CD16/32 antibody, followed by staining with primary antibodies: CD45 (I3/2.3) and CD326 (EpCAM) (G8.8) (BioLegend) for 20 minutes on ice. After sorting, isolated cells were pooled at the following ratios: CD45 EpCAM (keratinocytes), 5 parts; CD45 EpCAM (immune cells), 2.5 parts; and CD45 EpCAM (others), 2.5 parts. For the construction of single-cell RNA-Seq library, Chromium Next GEM Single Cell 3’ Library and Gel Bead Kit v3.1 (10x Genomics) was used according to the manufacturer’s instructions. Libraries were sequenced on a NovaSeq X plus (Illumina) at a read length of 28 x 90 to yield a minimum of 20,000 reads per cell for gene expression. The resulting raw data were processed by Cell Ranger 7.1.0 (10x Genomics). Further analysis was conducted using Seurat R package (version 4.2.1) with default parameters unless specified otherwise. For quality control purpose, we restricted the analysis to the cells (unique barcode) with exhibiting a percentage of mitochondrial gene < 5%, a total number of genes comprised between 200 and 5000. Data were normalized using the NormalizeData() function, and variable genes were identified using the FindVariableFeatures() function. Data scaling was performed with the ScaleData() function and principal components (PCs) were computed from these variable genes using the RunPCA() function. Clusters were identified using the FindNeighbors() and FindClusters() functions with a resolution parameter of 0.1. Non-linear dimensionality reduction and visualization were performed using UMAP via the RunUMAP() function. Clusters were categorized into 14 cell types based on marker genes: “keratinocytes”, “T cells”, “fibroblasts-1”, “fibroblasts-2”, “myeloid cells”, “hair follicle keratinocytes”, “dendritic cells-1”, “dendritic cells-2”, “ innate lymphoid cells”, “endothelial cells”, “sebocytes”, “mast cells”, “smooth muscle cells” and “others”. Further sub-clustering using the FindSubCluster() function with a resolution parameter of 0.1 resulted in the identification of four sub-clusters within the “hair follicle keratinocytes” and three sub-clusters within the “sebocytes.” This approach was used to create the final SeuratObject. Trajectory inference analysis was performed using Monocle3 (version 1.3.7). Seurat objects were converted into CellDataSet objects via the as.cell_data_set() function. Cells were embedded using UMAP, and trajectories were learned using the learn_graph() function. Pseudotime was assigned by specifying root cells (early time point). Cluster-wise pseudotemporal changes were visualized with plot_cells(). Genes whose expression changed along pseudotime were identified using Moran’s I statistic implemented in Monocle3. Genes with statistically significant Moran’s I values (adjusted *p* < 0.05) were considered to show pseudotime-dependent expression dynamics. These genes were subsequently subjected to KEGG pathway enrichment analysis using the clusterProfiler R package (version 4.14.6).

#### Statistical analysis

All analyses were performed using GraphPad Prism and R statistical language.^18^ Differences were considered significant when *p* < 0.05. The specific statistical tests are mentioned in Figure legends.

#### Illustration

Illustration was hand-drawn by the author using watercolor on paper. The illustration was minimally adjusted (color balance and cropping) using GIMP2.10.36 (GNU Image Manipulation Program). No AI-based image generation tools were used.

#### Language editing

The authors used ChatGPT (OpenAI) for language polishing, and the text was further revised by a native English speaker.

## Acknowledgments

We thank N Hoshino, Y Shirasaki, K Watanabe, M Baba, M Sakieda for technical assistance.

## References and Notes

### Funding

JSPS KAKENHI 20H03701 (YN) JST FOREST JPMJFR200Y (YN)

Joint Usage/Research Program of Medical Mycology Research Center, Chiba University. (YN and HT)

Research Grant on Inflammatory Skin Diseases, Japanese Dermatological Association (YN)

### Author contributions

Conceptualization: YN

Methodology: TS, NIS, YN,

Resources: TK, MM, YO, NI, HT, NIS, SY, MF, YN

Investigation: TS, MT, KT, AJH, SN, EI

Visualization: TS, MT, KT, AJH, NI, HT, NIS, YN

Illustration: YN

Funding acquisition: HT,YN,

Project administration: MF,YN

Supervision: NIS, SY, MF,YN

Writing – original draft: TS, NIS, YN

Writing – review & editing: BS, YN

All authors read and approved the final version of the submitted manuscript.

### Data and materials availability

All sequence data were deposited in the Sequence Read Archive (DNA Data Bank of Japan BioProject ID: PRJDB36936 and PRJDB37389).

## Inventory of upporting Information

### Supplementary data

**Table S1.** Correlation between cutaneous fatty acid levels and acne lesion counts. Spearman’s correlation coefficients (ρ, *Rs*) and p-values are shown for the relationship between individual cutaneous fatty acids (C14:0–C23:0, C16:1, C18:1) and the number of inflammatory or non-inflammatory comedones.

|  |  | C12:0 | C13:0 | C14:0 | C15:0 | C16:0 | 16:1(n-10) | C17:0 | C18:0 | C18:1(n-9) | C20:0 | C21:0 | C22:0 | C23:0 |
| --- | --- | --- | --- | --- | --- | --- | --- | --- | --- | --- | --- | --- | --- | --- |
| Inflammatory comedones | *Rs* | 0.20861 | 0.26609 | 0.31506 | 0.32875 | 0.29691 | 0.26164 | 0.31564 | 0.26328 | 0.27444 | 0.25317 | 0.080568 | 0.20609 | 0.22593 |
|  | *p value* | 0.146 | 0.061788 | 0.025843 | 0.019751 | 0.036271 | 0.066448 | 0.025556 | 0.064696 | 0.053771 | 0.076074 | 0.57808 | 0.15104 | 0.11465 |
| Non-inflammatory comedones | *Rs* | 0.1574 | 0.4349 | 0.1883 | 0.24647 | 0.1554 | 0.16843 | 0.24759 | 0.26022 | 0.22177 | 0.36664 | 0.31008 | 0.36854 | 0.41606 |
|  | *p value* | 0.27498 | 0.0015987 | 0.19034 | 0.084445 | 0.28119 | 0.24232 | 0.083001 | 0.067987 | 0.12166 | 0.0088217 | 0.028418 | 0.008452 | 0.0026545 |
| Total comedones | *Rs* | 0.31718 | 0.51969 | 0.41841 | 0.45613 | 0.38381 | 0.35875 | 0.44439 | 0.36264 | 0.41868 | 0.41177 | 0.2258 | 0.36861 | 0.37395 |
|  | *p value* | 0.024806 | 0.0001099 | 0.002496 | 0.0008712 | 0.0059307 | 0.010517 | 0.0012247 | 0.0096483 | 0.0024784 | 0.0029679 | 0.11486 | 0.0084384 | 0.007469 |

**Figure S1.**
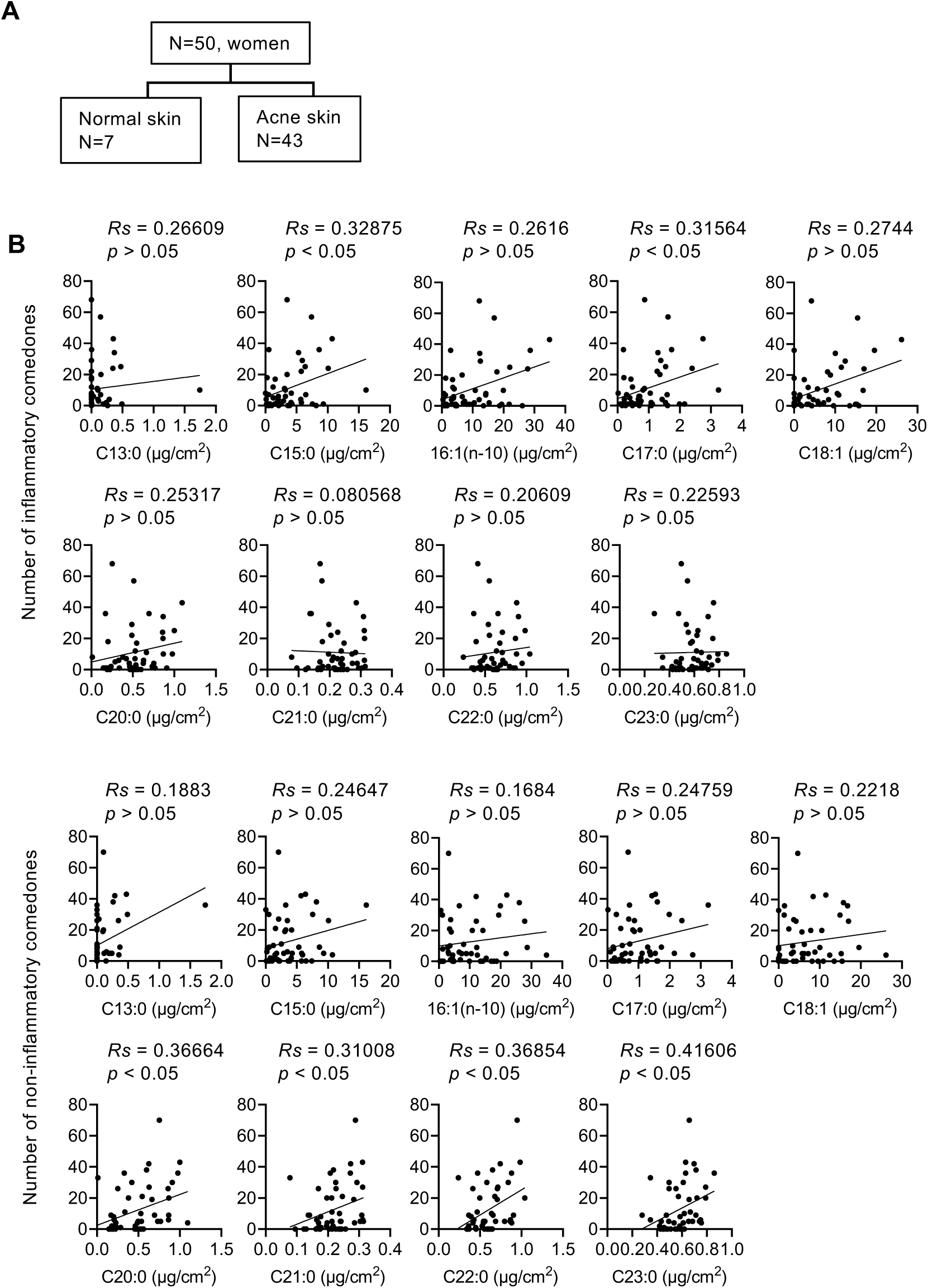
Study design and correlation analysis of cutaneous lipids. **(A)** Schematic representation of the study workflow. A total of 50 Japanese female participants with either normal skin (N = 7) or acne-prone skin (N = 43) were recruited. **(B)** Correlation analysis between cutaneous fatty acid levels and clinical acne lesion counts, including both inflammatory and non-inflammatory comedones.

**Figure S2.**
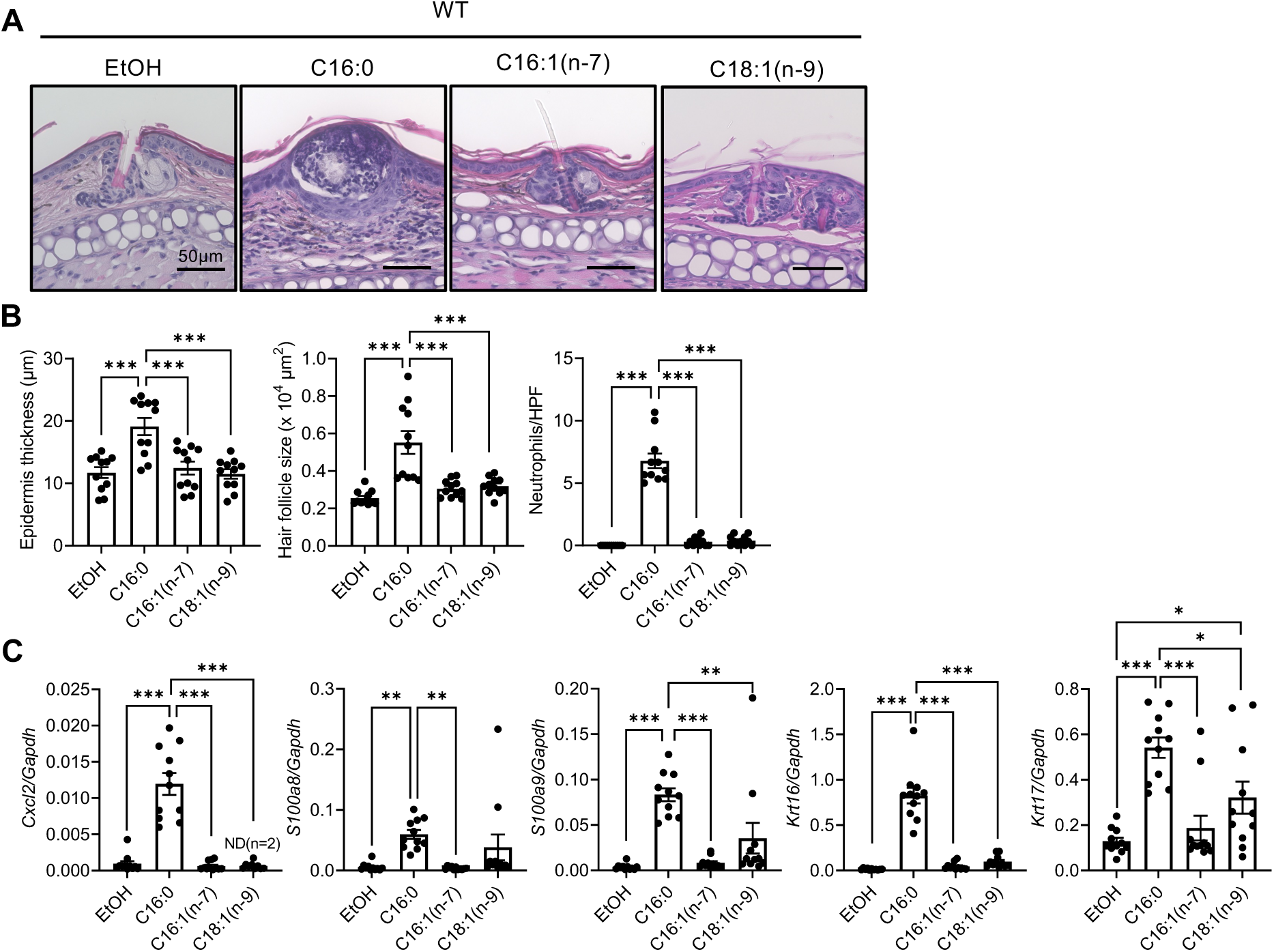
C16:0 uniquely induces follicular inflammation and keratinocyte activation compared to unsaturated fatty acids. **(A)** Representative H&E-stained sections of mouse ear skin two days after topical application of C16:0, C16:1, or C18:1. **(B)** Measurements of ear thickness, follicle size, and inflammatory cell infiltration in mice two days after topical application of FAEEs ranging from C14:0 to C18:0. **(C)** Relative mRNA expression levels of *Cxcl2, S100a8, S100a9, Krt16* and *Krt17* in whole ear skin following treatment. Data points represent individual mice and bars represent mean ± SEM. All data were collected from at least two independent experiments. Controls refer to vehicle-treated skin (EtOH, ethanol). ∗p < 0.05; ∗∗p < 0.01; ∗∗∗p < 0.001. Data were analyzed by one-way ANOVA with Dunnett’s T3 multiple comparison test.

**Figure S3.**
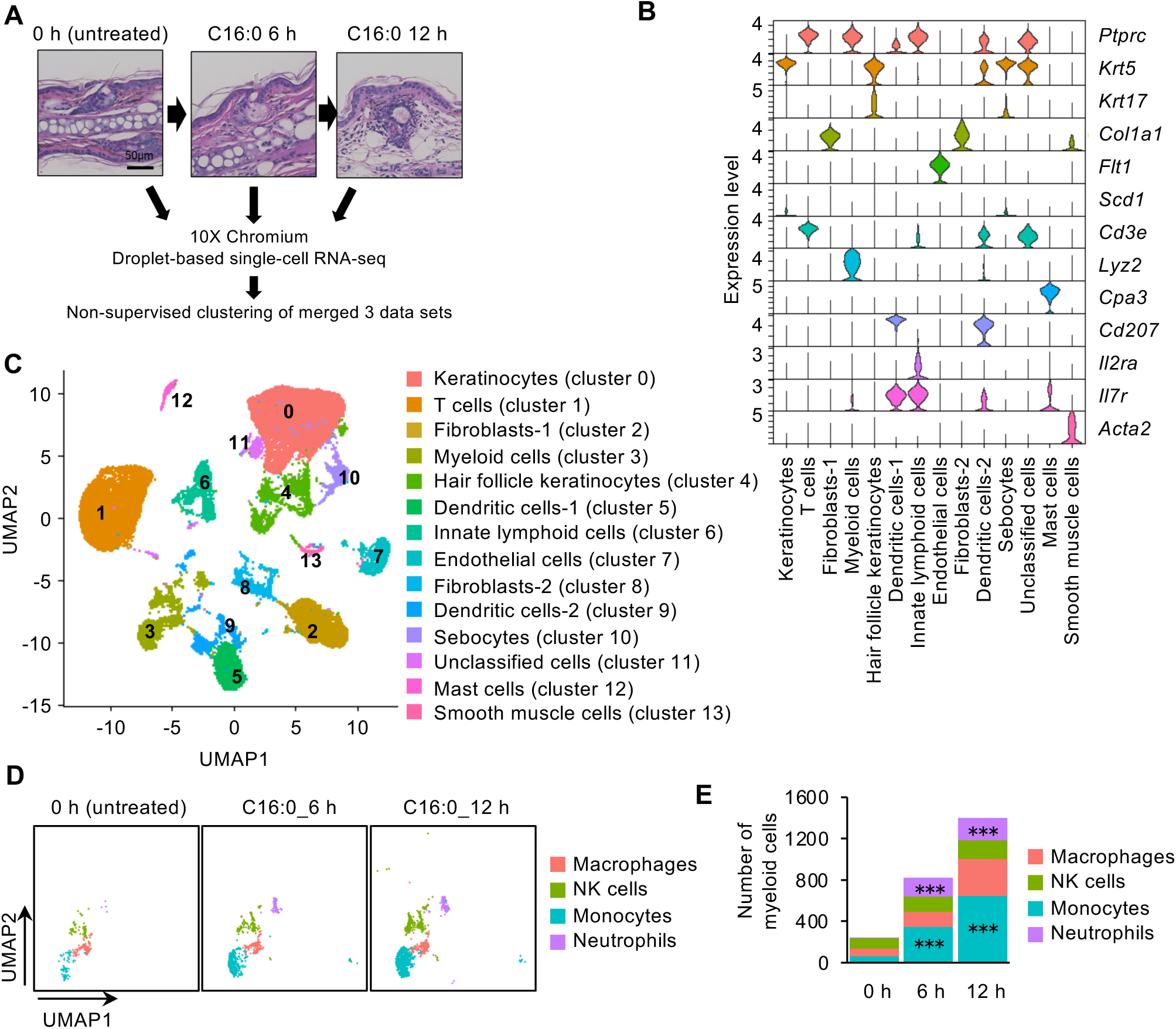
Single-cell transcriptomic analysis reveals C16:0 induced expansion of inflammatory myeloid populations in the skin. **(A)** Experimental timeline and single-cell RNA sequencing (scRNA-seq) design. Ear skin was collected from untreated (0 hour,0 h) control and C16:0-treated mice at 6 (6 h) and 12 hours (12 h) post-application. **(B)** Expression levels (y axis) of cluster-defining genes in each cluster. Violin plots show the distribution of the normalized expression levels of genes. **(C)** UMAP plot displaying 14 transcriptionally distinct cell populations (clusters 0-13), annotated based on marker gene expression and representing major skin-resident cell types. **(D, E)** Quantification of immune cell subsets within the myeloid compartment (cluster 3) derived from scRNA-seq data. Data are from a single experiment with five biological replicates (individual mice). Statistical significance in (E) was assessed by chi-square test followed by analysis of standardized residuals to identify cell populations contributing to overall distribution changes. ∗p < 0.05; ∗∗p < 0.01; ∗∗∗p < 0.001.

**Figure S4.**
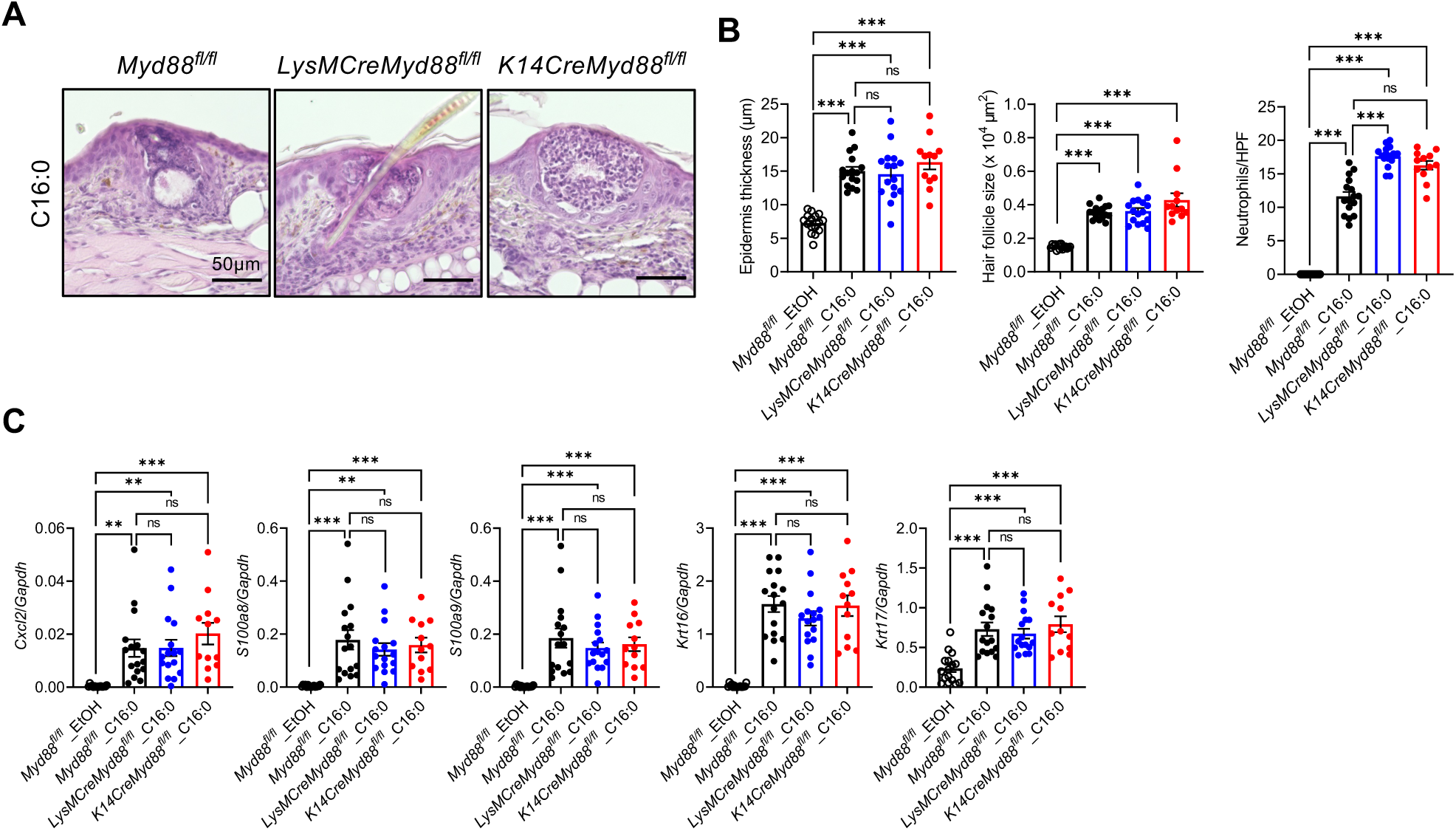
Keratinocyte- and myeloid-specific Myd88 deletion does not affect C16:0 induced skin responses. **(A)** Representative H&E-stained sections of mouse ear skin two days after topical application of C16:0 in *Myd88^f/f^* control, *K14CreMyd88^f/f^* (keratinocyte-specific knockout), and *LysMCreMyd88^f/f^* (myeloid-specific knockout) mice. Scale bars, 50 μm. **(B)** Measurements of ear thickness, follicle size, and inflammatory cell infiltration in the same groups as in (A). *Myd88^f/f^* **(C)** Gene expression analysis *of Cxcl2, S100a8*, *S100a9, Krt16 and Krt17* in whole ear skin following C16:0 treatment. Data points represent individual mice and bars represent mean ± SEM. All data were collected from at least two independent experiments. Controls refer to vehicle-treated skin (EtOH, ethanol). ∗p < 0.05; ∗∗p < 0.01; ∗∗∗p < 0.001; ns, not significant. Data were analyzed by one-way ANOVA with Dunnett’s T3 multiple comparison test.

**Figure S5.**
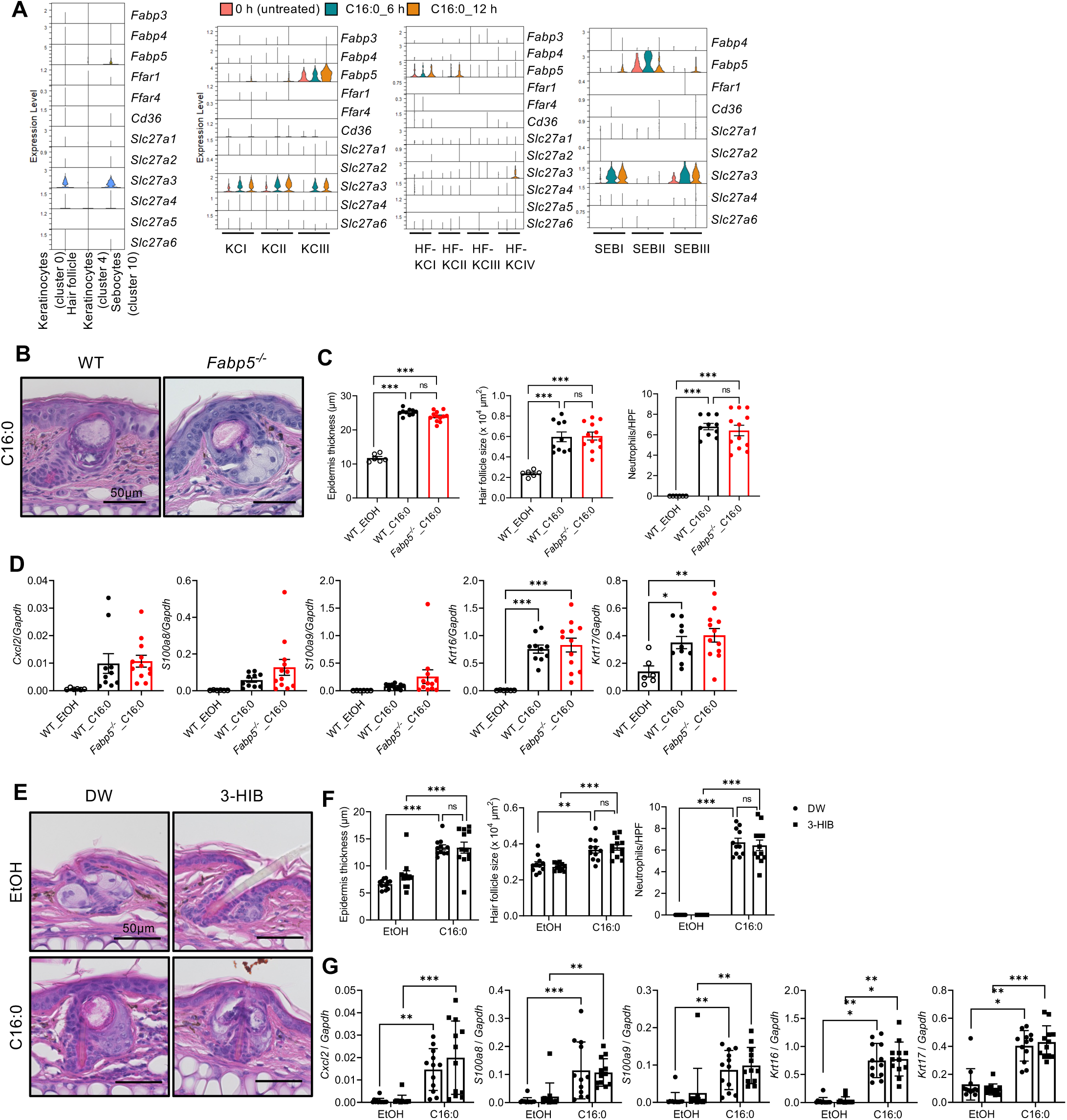
Fatty acid transporter activity is not required for C16:0 induced inflammation. **(A)** Expression levels of fatty acid transporters (*Fabp3*, *Fabp4*, *Fabp5, Cd36*, *Slc27a1*, *Slc27a2*, *Slc27a3* (*Fatp3*), *Slc27a4*, *Slc27a6*) in *Lrig1^+^* progenitor cells from the scRNA-seq dataset. **(B, C)** Representative histological images (B) and measurements of ear thickness, follicle size, and inflammatory cell infiltration (C). Scale bars, 50 μm. **(D)** Quantification of inflammatory gene expression (*Cxcl2*, *S100a8, S100a9*) and keratinization markers (*Krt16, Krt17*) in the ears of *Fabp5* knockout (KO) and wild-type (WT) mice two days after topical application of C16:0. **(E, F)** Representative histological images (E) and measurements of ear thickness, follicle size, and inflammatory cell infiltration (F). Scale bars, 50 μm. **(G)** Expression of inflammatory genes (*Cxcl2*, *S100a8, S100a9*) and keratinization markers (*Krt16, Krt17*) in ear skin following 3-HIB and C16:0 treatment. Data in (A) are from a single experiment with five biological replicates (individual mice). Data points in (C, D, F, G) represent individual mice; bars indicate mean ± SEM. Data in (B)-(G) are from at least two independent experiments. Controls refer to vehicle-treated skin (EtOH, ethanol). ∗p < 0.05; ∗∗p < 0.01; ∗∗∗p < 0.001; ns, not significant. Data were analyzed by one-way ANOVA with Dunnett’s T3 multiple comparisons test.

